# Mugs and Plants: Object Semantic Knowledge Alters Perceptual Processing with Behavioral Ramifications

**DOI:** 10.1101/2021.03.24.436853

**Authors:** Dick Dubbelde, Sarah Shomstein

## Abstract

Neural processing of objects with action associations recruits dorsal visual regions more than objects without such associations. We hypothesized that because the dorsal and ventral visual pathways have differing proportions of magno- and parvo-cellular input, there should be behavioral differences in perceptual tasks between manipulable and non-manipulable objects. This hypothesis was tested in adults across five experiments (Ns = 26, 26, 30, 25, 25) using a gap detection task, suited to the spatial resolution of parvocellular processing, and an object flicker discrimination task, suited to the temporal resolution of magnocellular processing. Directly predicted from the cellular composition of each pathway, a strong non-manipulable object advantage was observed in gap detection, and a small manipulable object advantage in flicker discrimination. Additionally, these effects are modulated by reducing object recognition through inversion and by suppressing magnocellular processing using red light. These results establish perceptual differences between objects dependent on semantic knowledge.

**Statement of Relevance:** When we perceive an object, our knowledge of that object is brought to mind. Previous work has shown specifically that knowledge of object manipulability biases neural processing to areas of the brain in the parietal lobe which are relevant to motor processing. In this study we show that this neural bias, caused by knowledge of the object, has an effect on object perception. Using behavioral paradigms designed to take advantage of the specific response properties of neurons in the parietal and temporal object processing areas, we found that manipulable objects are perceived with higher temporal resolution while non-manipulable objects are perceived with higher spatial resolution. Our results demonstrate a specific neural mechanism by which prior knowledge affects current perception.

## Introduction

Object perception underlies effective engagement with our environment. When examining a potted plant, for example, precise access to long term semantic memory may be needed to facilitate detailed identification of the plant to determine if it needs water. However, when reaching for a mug with your morning coffee, detailed identification may be less important (e.g., which one of your 10 favorite mugs it is), but computing a motor plan to bring the mug to your mouth without spilling is pertinent. The visual object processing necessary to identify plants and manipulate mugs progresses along two distinct but interacting neural processing pathways, each of which subserves different end goals (Mishkin & Ungerleider, 1982; Kravitz et al., 2011). While the hypothesis of biased processing across the two pathways is widely accepted, it is unclear whether there are behavioral ramifications that directly relate to object processing being biased to different pathways. We provide evidence that object semantic knowledge evokes differences in object perception based on the recruited regions across the visual pathways. Importantly, we gain insight into neural processing by using purely behavioral methods that are custom tailored to the response properties of the neurons in both the dorsal (higher temporal resolution) and ventral pathways (higher spatial resolution).

The ventral pathway, known as the ‘what’ pathway, is characterized by object feature selectivity and projects anteriorly from the occipital visual areas toward the hippocampus and the inferior temporal cortex (Kravitz et al., 2013). The dorsal pathway, colloquially known as the ‘where’ or ‘how’ pathway, courses from the occipital visual areas into the parietal cortex. The dorsal pathway is understood to subserve object processing for the purposes of reaching, grasping, and manipulation. While both pathways contribute to visual perception, the demands of perceiving a specific object can differentially engage the two pathways such that the dorsal pathway provides greater contribution for visual processing of objects that are directly relevant for manipulation (e.g., hammer, plunger, or mug). Evidence for dynamic and biased recruitment of the visual pathways has been garnered from a wide range of techniques and paradigms. For example, while presenting participants with objects of various categories, including tools, places, animals, and faces, an increase in dorsal pathway processing, specifically in the left ventral premotor and posterior parietal cortices, is observed exclusively in response to tool presentation (Chao & Martin, 2000). Similar results evidencing a dorsal bias for manipulable objects are observed in functional magnetic resonance imaging studies (Noppeney et al., 2006a; Mahon et al., 2007, Almeida et al., 2013, Chen et al., 2018) and with various other paradigms including interocular suppression (Fang and He, 2005) and continuous flash suppression (Almeida et al., 2008; Almeida et al., 2010; but see Sakuraba et al., 2012; Almeida et al., 2014).

While neurophysiological evidence has been convincing in showing selectivity for objects across the two streams, the impact that differential processing across the two pathways has on object perception and subsequent behavior has not been characterized. Hypotheses regarding how dynamic recruitment of the visual pathways influences perception and behavior are based on an asymmetry in the cellular innervation of the two pathways that overlaps with the separate magnocellular and parvocellular channels identified in anatomical architecture of visual processing (Maunsell et al., 1990; Baizer et al., 1991; Ferrera et al., 1992). The asymmetry in input endows each pathway with different response properties in accordance with cell stimulus preferences within the magno- and parvo-cellular channels. The magno- and parvo-cellular channels originate from different types of ganglion cells within the retina and course separately through different layers of the lateral geniculate nucleus (Leventhal et al., 1981; Perry et al., 1984) to innervate separate layers of V1 (Blasdel & Lund, 1983). From V1, the parvocellular channel can be followed into V2, V4, and then into the inferior parietal cortex while the magnocellular channel can be traced into different regions of V2, through V3d and MT, and into the posterior parietal cortices (DeYoe & van Essen, 1988; Livingstone & Hubel, 1988).

Differing innervation of the two visual pathways leads to different response properties that convey information of different spatial and temporal resolutions. The heavily myelinated magnocellular channel is derived from the parasol ganglion cells with relatively large receptive fields, spanning large regions of the retina (Maunsell et al., 1999a; Nassi & Callaway, 2009). The features of the parasol retinal ganglion cells enable the dorsally biased magnocellular channel to encode information with higher temporal resolution than the parvocellular channel (Pokorny and Smith, 1997; but see Maunsell et al., 1999b). Conversely, the parvocellular channel is derived from the midget retinal ganglion cells which receive input primarily from cone receptors and have smaller receptive fields (Nassi & Callaway, 2009). The features of the midget retinal ganglion cells enable the ventrally biased parvocellular channel to encode information with a higher spatial resolution than the magnocellular channel (Derrington & Lennie, 1984; Leonova et al., 2003; McAnany & Alexander, 2008).

The connections between the magnocellular channel and temporal resolution and the parvocellular channel and spatial resolution have been used to study the neural basis of several behavioral effects ranging from spatial attention (Yeshurun, 2004) to the effects of fear on visual perception (Bocanegra and Zeelenberg, 2009). Most relevantly, the asymmetric contributions of the magno- and parvo-cellular channels to the dorsal and ventral pathways have been proposed as a mechanism for near-hand effects on perception (Gozli et al., 2012; Chan et al., 2013). In one such study, it was hypothesized that when the participants’ hands were near the stimuli then processing would be biased to the dorsal pathway, which is more predominantly magnocellular, leading to enhanced sensitivity to temporal blinks. Conversely, when the participants’ hands were far from the stimuli then processing would be biased to the ventral pathway, which is more predominantly parvocellular, leading to an enhanced sensitivity to spatial gaps (Gozli et al., 2012). Consistent with differential engagement of two streams, it was observed that placing hands near the stimuli increased participants’ sensitivity to temporal blinks, while moving hands away from the screen increased sensitivity to spatial gaps.

Knowing that semantic knowledge of manipulability biases processing to the dorsal stream, and that the dorsal and ventral streams have different response profiles due to differential innervation by the magnocellular and parvocellular channels, we hypothesize that semantic knowledge of object manipulability should have consequences for perceptual processing. Thus, because semantic knowledge of an object’s manipulability determines which pathway will be biased, we predict that (1) manipulable objects, that elicit a higher degree of magnocellularly biased dorsal processing, are processed with higher temporal resolution and (2) non-manipulable objects, that rely more on parvocellularly biased ventral processing, are processed with higher spatial resolution. Across five experiments, these hypotheses are tested by comparing spatial and temporal resolution across two object groups: manipulable and non-manipulable objects.

## Method

### Participants

All participants were recruited from The George Washington University participant pool, gave informed consent according to the George Washington University’s institutional review board (IRB), were naïve to the purpose of the experiment, and all reported normal or corrected- to-normal vision. In Experiment 4, which utilized color stimuli no participant reported colorblindness.

For each experiment, sample sizes were chosen based on behavioral studies demonstrating similar effects (i.e., Gozli et al., 2012) and participants were recruited in batches of 6-10 until at least 25 participants with accuracy above chance were accumulated. For Experiment 1, 26 undergraduate students (19 female; average age: 19.2; 6 left-handed) were recruited. No participant was removed from the analyses. For Experiment 2, 27 undergraduate students were recruited. Twenty-six participants’ data (14 female; average age: 19.0; 1 left-handed) were analyzed after one participant was cut for below chance accuracy in one of the conditions. For Experiment 3, 33 undergraduate students were recruited. Thirty participants’ data (26 female; average age: 19.07, 3 left-handed) were analyzed after three participants were cut for below chance accuracy in at least one of the conditions. For Experiment 4, 40 undergraduate students were recruited. Twenty-five participants’ data (14 female; average age: 19.0; 1 left-handed) were analyzed after six participants were cut for not finishing the experiment and nine participants were cut for having less than chance accuracy in at least one of the conditions. For Experiment 5, 26 undergraduate students were recruited. Twenty-five participants’ data (18 female; average age: 19.5, 0 left-handed) were analyzed after one participant was cut for chance accuracy in all conditions.

### Apparatus and Stimuli

All experiments were presented on a 24” Acer GN246 HL monitor with a refresh rate of 144 Hz, positioned at a distance of 60 cm from the viewer in a dark room. The experiment was generated and presented using PsychoPy v1.82.

The stimuli in Experiments 1, 2, 4, and 5 consisted of line drawings of real-world, everyday objects drawn both from The Noun Project, an online repository of object icons and clip art (https://thenounproject.com). The object stimuli consisted of 10 line-drawings of manipulable objects, and 10 line-drawings of non-manipulable objects. The manipulable objects were as follows: snow shovel, handsaw, plunger, screwdriver, hammer, wrench, knife, bottle opener, spatula, and mug. The non-manipulable objects were: fire hydrant, picture frame, window, toilet, candle, garbage can, water fountain, potted plant, fan, and lamp. The stimuli were displayed as large as possible in a 4° x 4° area and all stimuli were presented in black (HSV = 0, 0, 0) on a dark gray background (HSV = 0, 0, 50). All objects are displayed in Figure 1. Objects were controlled for low-level differences by calculating the mean luminance, size, aspect ratio (i.e., measure of elongation), and spatial frequency (average distance from origin of the points calculated from a 2D fast Fourier transform) for each object. Assessed with independent samples t-tests, there were no mean differences between object groups in luminance(t(18) = -0.960, p = 0.350), mean size (t-test (t(18) = 1.043, p = 0.311), aspect ratio (t(18) = 1.209, p = 0.242), or spatial frequency (t-test (t(18) = -0.155, p = 0.879). The width of the bottom line was controlled for each object on which the gap appeared. There was no significant difference between manipulable and non-manipulable objects in bottom line width in an independent samples t-test (t(18) = 0.922, p = 0.369).

**Figure 1.**
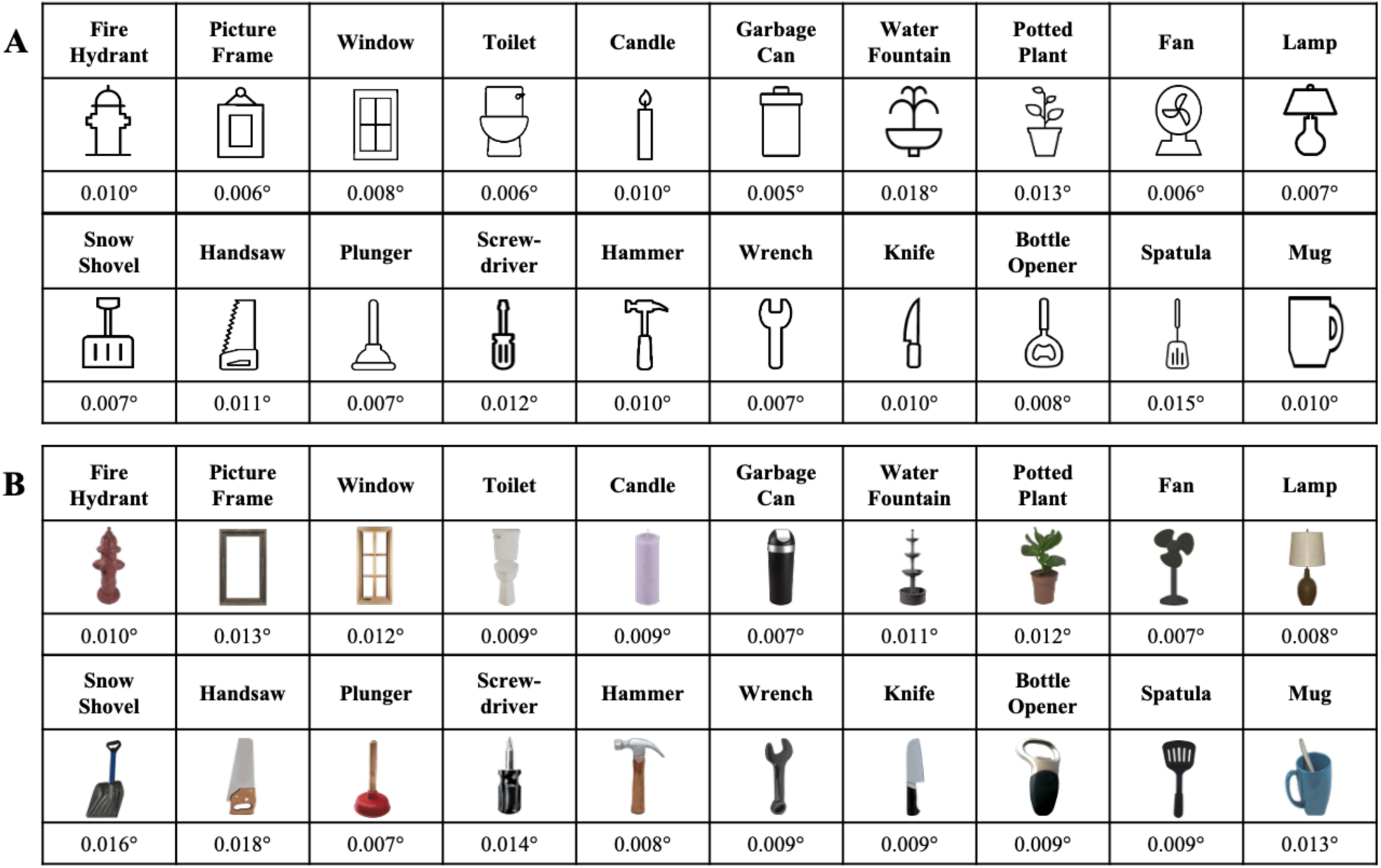
Stimulus set used in the experimental paradigms. (A) Line drawings of manipulable and non-manipulable objects used in Exp. 1. (B) Pictures of real-world manipulable and non-manipulable objects used in Experiment 3. The numbers beneath each set of objects correspond to the size of the spatial gap titrated to each object.

For Experiment 3, the stimuli consisted of images of real-world, everyday objects drawn from Google image search and manipulated in GIMP 2.10.4. The object stimuli consisted of 10 images of the same manipulable objects, and 10 images of the same non-manipulable objects used in Experiments 1,2, 4, and 5. The stimuli were displayed as large as possible in a 4° x 4° area and all stimuli were presented in color on a dark gray background (HSV = 0, 0, 50). All objects are displayed in Figure 1.

Additionally, the colored objects used in Experiment 3 were similarly controlled for low-level differences by calculating the mean size, hue, saturation, value, aspect ratio, and spatial frequency. There was no significant differences between manipulable and non-manipulable objects in an independent samples t-test for mean size (t(18) = -1.027, p = 0.318)hue (t(18) = -0.309, p = 0.761), saturation (t(18) = -0.031, p = 0.976), value (t(18) = 0.032, p = 0.975), aspect ratio (t(18) = -0.094, p = 0.927), or spatial frequency (t(18) = -1.466, p = 0.160) Additionally, we controlled the width of the bottom line of each object on which the gap would appear. There was no significant difference between manipulable and non-manipulable objects in bottom line width in an independent samples t-test (t(18) = 0.557, p = 0.584).

### Task

In Experiment 1, each trial began with a single central fixation point, which subtended a visual angle of 1° x 1°. The central fixation point was rendered in white (HSV = 0, 0, 100). After 1000 ms of fixation presentation, a single object line drawing would appear 4° to the left or right of the fixation point. The side of presentation was counterbalanced such that each participant saw an equal number of stimuli on the left and on the right. The stimuli were displayed as large as possible within a 4° x 4° area and all stimuli were presented in color on a dark gray background (HSV = 0, 0, 50). All objects are displayed in Figure 1.

In one half of the experiment, the object appeared with or without a spatial gap in the center of the bottom line of the object. The objects had an equal probability of having a spatial gap or not having a spatial gap. The objects appeared for 100 ms and participants were to report the presence of the spatial gap by pressing the right control button (present) or the left control button (absent) on the keyboard. Feedback was presented on incorrect trials only.

A staircase procedure was used to calibrate the size of the gap to each object to ensure that the gap was equally perceptible across each object regardless of each objects’ individual characteristics (Pelli and Bex, 2013). Sixteen undergraduate students (13 female; average age: 18.8; 4 left-handed) from the George Washington University participated in exchange for course credit. In each trial the object would be presented with or without the spatial gap which would begin with a size of 0.025° visual angle. If the gap was detected correctly for two consecutive trials, the gap decreased by one 0.005° step. If the gap was missed, or a false alarm was made to the absence of a gap, for two consecutive trials, the gap was increased by one 0.005° step. The gap was calibrated until twenty trials had been completed or until the staircase reversed direction (2 correct followed by 2 incorrect or vice versa) three times. The gap size for each object is displayed in figure 1.

In the other half of the experiment, the object appeared with or without a temporal blink. The objects had an equal probability of having a temporal blink or not having a temporal blink. The blinks were 16 ms long. The object first appeared for 96 ms, followed by the temporal blink, and the object then appeared a second time for 32 ms. Participants were to report the presence of the temporal blink by pressing the right control button and absence of the temporal blink by pressing the left control button on the keyboard. Feedback was presented on incorrect trials only.

The presentation of stimulus type was counterbalanced across participants such that half of the participants were first presented with spatial gaps and half of the participants were presented with temporal blinks first. Participants completed a total of 720 trials broken into 6 blocks, 360 trials of spatial gaps and 360 trials of temporal blinks.

Experiment 2 was identical to Experiment 1 except each stimulus was presented upside down. The spatial gap was presented in the same place as it was in Experiment 1 relative to the object.

Experiment 3 was identical to Experiment 1, except more realistic images were used instead of line drawings. Similarly to the line drawing stimuli, a staircase procedure was used to calibrate the size of the gap to each object to ensure that the gap would be equally perceptible across each object regardless of each objects’ individual characteristics (Pelli and Bex, 2013). Twenty-three undergraduate students (18 female; average age: 18.9; 4 left-handed) from the George Washington University participated in exchange for course credit and the same procedure from Experiment 1 was used. The gap size for each object is displayed in figure 1.

Experiment 4 was similar in trial structure to the temporal blink condition of Experiment 1, except instead of temporal blink detection each trial had a temporal blink that was either short or long in duration. The objects had an equal probability of having a short or a long temporal blink. In the short blink condition, the object first appeared for 96 ms, followed by a 16 ms temporal blink, and then the object appeared a second time for 32 ms. In the long blink condition, the object first appeared for 64 ms, followed by a 48 ms temporal blink, and then the object was presented for a second time for 32 ms. Participants were to report a presence of a short temporal blink by pressing the ‘c’ key and a long temporal blink by pressing the ‘m’ key on the keyboard. Feedback was presented exclusively on incorrect trials. Participants completed a total of 720 trials broken into 6 blocks, 360 trials of short blink duration and 360 trials of long blink duration.

Experiment 5 was identical to the spatial gap procedure of Experiment 1 except the background color was either green (HSV = 110, 30, 90) or red (HSV = 5, 30, 90). The background color was counterbalanced across participants such that half of the participants would first be presented with the green background and half of the participants would first be presented with the red background. Participants completed a total of 720 trials broken into 6 blocks, 360 trials of green backgrounds and 360 trials of red backgrounds.

## Results

### Experiment 1

The spatial and temporal paradigms used d’ as a measure of perceptual sensitivity (Fig. 2). A two-way repeated-measures analysis of variance (ANOVA) was conducted on d’ with object group (manipulable, non-manipulable) and stimulus type (gap, blink) as within-subject variables (Fig. 2C, left). The ANOVA revealed no significant main effect of object group (*F*(1, 25) = 2.342, *p* = 0.138, *η*_*p*_^*2*^ = 0.086), and a significant main effect of task type (*F*(1, 25) = 50.953, *p* < 0.001, *η*_*p*_^*2*^ = 0.671) such that d’ sensitivity was higher for blinks than for gaps (M_gaps_= 1.926, 95% CI [1.673, 2.179]; M_blinks_ = 2.967, 95% CI [2.680, 3.254]). Importantly, and consistent with the hypothesis of differential engagement of two pathways depending on object utility, a significant two-way interaction between object group and task type (*F*(1, 25) = 6.772, *p* = 0.015, *η*_*p*_^*2*^ = 0.213) was observed, driven by the difference between object groups in the gap condition (*F*(1, 25) = 9.888, *p* = 0.004, *η*_*p*_^*2*^ = 0.283), with non-manipulable objects having a higher d’ for gaps than manipulable objects (M_non-manipulable_ = 2.021, 95% CI [1.756, 2.285]; M_manipulable_ = 1.831, 95% CI [1.589, 2.073]). The interaction effect and the driving simple main effect are consistent with the prediction that non-manipulable objects, given their higher reliance on the ventral pathway, should yield higher sensitivity in the detection of spatial gaps than manipulable objects. Notably, the expected higher sensitivity to temporal gaps in the manipulable object set was not supported. The possibility of an advantage for manipulable objects in temporal sensitivity is further examined in Experiment 4.

**Figure 2.**
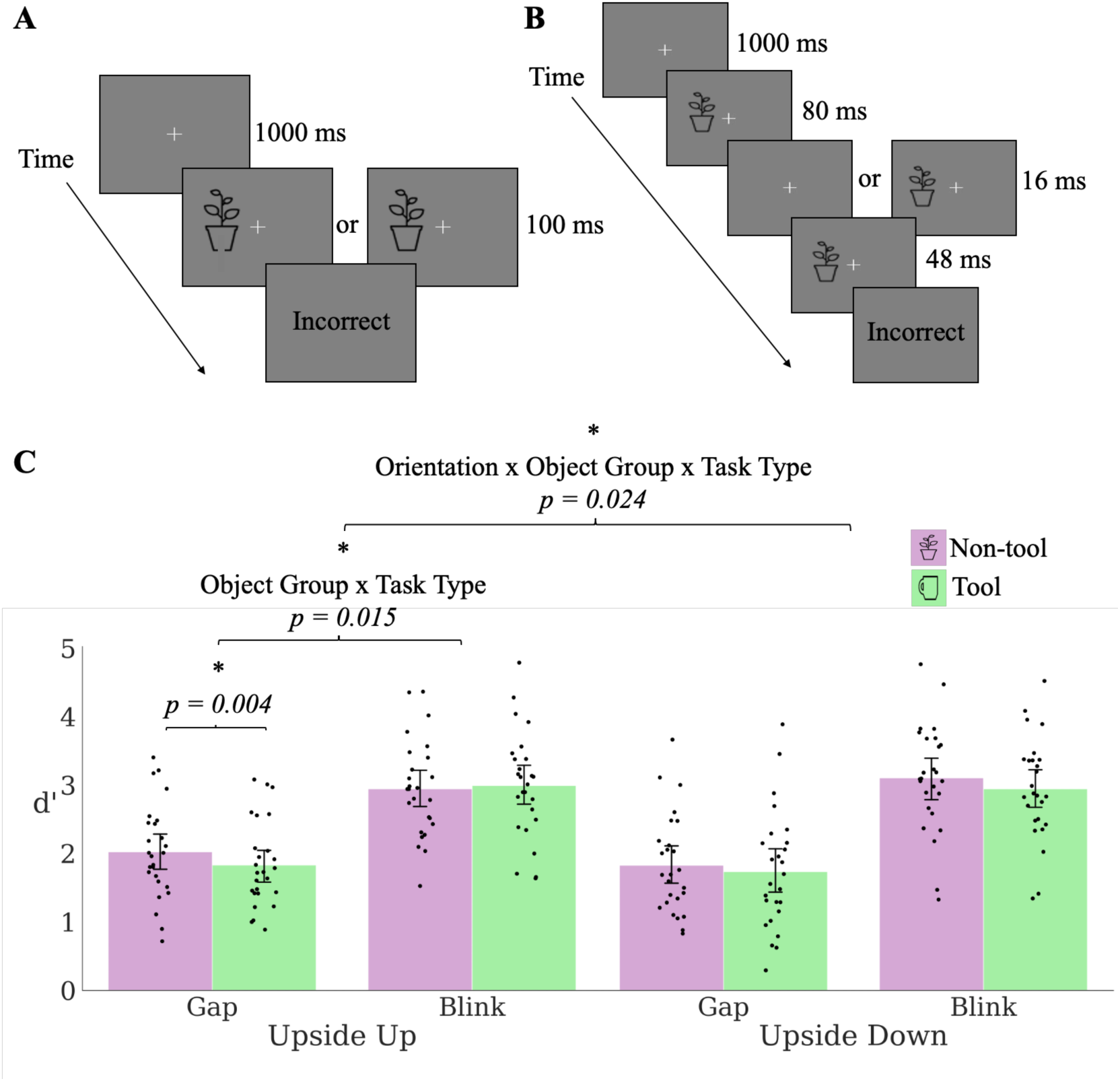
The paradigm and results for Experiments 1 and 2. Participants maintained fixation on the center cross. An object appeared to the left or right of fixation. A) For the spatial-gap paradigm, participants indicated by keypress if the bottom line of the presented object contained a gap. B) For the temporal paradigm, participants were to indicate whether the object flickered. C, left) The results from Experiment 1 in which the object stimuli were presented upside up. C, right) The results from Experiment 2 in which the objects were presented upside down.

### Experiment 2

To further probe the hypothesis that semantic knowledge of object utility (manipulable or non-manipulable) biases the pathway that will ultimately process the object, we designed Experiment 2 to demonstrate that the hypothesized pathway biasing will not occur if the semantic identity of the objects is obscured (Firestone and Scholl, 2016). The same paradigm and objects from Experiment 1 were used but the objects were inverted (upside-down). The inversion preserved the low-level features of each object while impairing participants’ ability to rapidly recognize and access their semantic knowledge of the objects’ function. Inversion has been found to interfere with recognition of faces and objects (Diamond and Carey, 1986).

Following from the hypothesis that semantic knowledge of object’s utility drives the perceptual difference between manipulable and non-manipulable objects, we predict that object inversion should reduce the difference between manipulable and non-manipulable objects. A two-way repeated-measures ANOVA was conducted on d’ with object group (manipulable, non-manipulable) and task type (gap, blink) as within-subject variables. The ANOVA revealed a significant main effect of object group (*F*(1, 25) = 5.860, *p* = 0.023, *η*_*p*_^*2*^ = 0.190) such that d’ was higher for non-manipulable objects than for manipulable objects (M_non-manipulable_ = 2.466, 95% CI [2.174, 2.758]; M_manipulable_ = 2.340, 95% CI [2.029, 2.650]) (Fig. 2C, right). There was also a significant main effect of task type (*F*(1, 25) = 82.298, *p* < 0.001, *η*_*p*_^*2*^ = 0.767) such that d’ was higher for blinks than for gaps (M_gaps_ = 1.782, 95% CI [1.476, 2.088]; M_blinks_ = 3.023, 95% CI [2.723, 3.320]). As predicted, there was no significant interaction between object group and task type (*F*(1, 25) = 0.508, *p* = 0.483, *η*_*p*_^*2*^ = 0.020).

To assess the non-significant interaction effect, a Bayes factor analysis was conducted using the Bayesian repeated measures ANOVA in JASP (van den Bergh et al., 2019) comparing the posterior probability of a model with the main effects of stimulus type and object group but no interaction term as the null hypothesis (*H*_0_) to the posterior probability of a full model with the main effects and the interaction term as the alternative hypothesis (*H*_1_). The Bayes factor analysis evaluating the non-significant interaction effect yielded a Bayes factor of 4.83 (*BF*_01_=4.83), indicating substantial evidence for the null hypothesis which does not include the interaction (Kass & Raftery, 1995).

Lastly, in order to statistically demonstrate that results of the inversion experiment are indeed different from those observed in Experiment 1, data were subjected to a between-subject ANOVA with object orientation (upright, inverted) as a between subjects variable and object group and task type as within subject factors. The ANOVA revealed a significant three-way interaction between object group, task type, and orientation (*F*(1, 50) = 5.381, *p* = 0.024, *η*_*p*_^*2*^ = 0.097) such that, as predicted, orientation significantly reduced the effect for the inverted objects.

### Experiment 3

While Experiment 1 provides evidence for biased engagement of the ventral pathway for non-manipulable object perception in a spatial task, it could be argued that despite careful low-level feature controls (e.g., luminance, size, elongation, spatial frequency) an uncontrolled low-level difference between manipulable and non-manipulable objects is responsible for driving the manipulable vs. non-manipulable advantage in the spatial gap task.

Experiment 3 used the same paradigm as Experiment 1 except the line drawings were replaced by real-world images of corresponding objects, such that a line drawing of a candle was replaced with a picture of a candle, etc. (Fig. 1B). The prediction remained the same as in the original experiment. If object semantic knowledge determines which visual pathway object processing is biased to, then non-manipulable objects will be biased to the ventral pathway, leading to higher sensitivity (as measured by d’) in the spatial gap task. This would replicate the pattern of performance observed in Experiment 1. A two-way repeated-measures ANOVA was conducted on d’ with object group (manipulable, non-manipulable) and task type (gap, blink) as within-subject variables. The ANOVA revealed a significant main effect of object group (*F*(1, 29) = 28.211, *p* < 0.001, *η*_*p*_^*2*^ = 0.493) with non-manipulable objects having a higher sensitivity than manipulable objects (M_non-manipulable_ = 2.361, 95% CI [2.054, 2.668]; M_manipulable_ = 2.047, 95% CI [1.773, 2.320]) (Fig. 3). There was also a significant main effect of task type (*F*(1, 29) = 8.413, *p* = 0.007, *η*_*p*_^*2*^ = 0.225) with blinks having a higher average sensitivity than gaps (M_gaps_ = 1.998, 95% CI [1.758, 2.238]; M_blinks_ = 2.410, 95% CI [2.068, 2.751]). Importantly, a two-way interaction between object group and task type was also significant (*F*(1, 29) = 13.061, *p* = 0.001, *η*_*p*_^*2*^ = 0.311), driven by a simple main effect in the gap condition (*F*(1, 29) = 46.424, *p* < 0.001, *η*_*p*_^*2*^ = 0.616) such that non-manipulable objects had a higher d’ for gaps than did the manipulable objects (M_non-manipulable_ = 2.238, 95% CI [1.982, 2.494]; M_manipulable_ = 1.758, 95% CI [1.535, 1.981]). In addition to serving as a low-level control and a further test of the hypothesis, these results replicate Experiment 1, providing strong additional support for the hypothesis that the perceptual differences between manipulable objects and non-manipulable objects is due to the semantic content of the objects.

**Figure 3.**
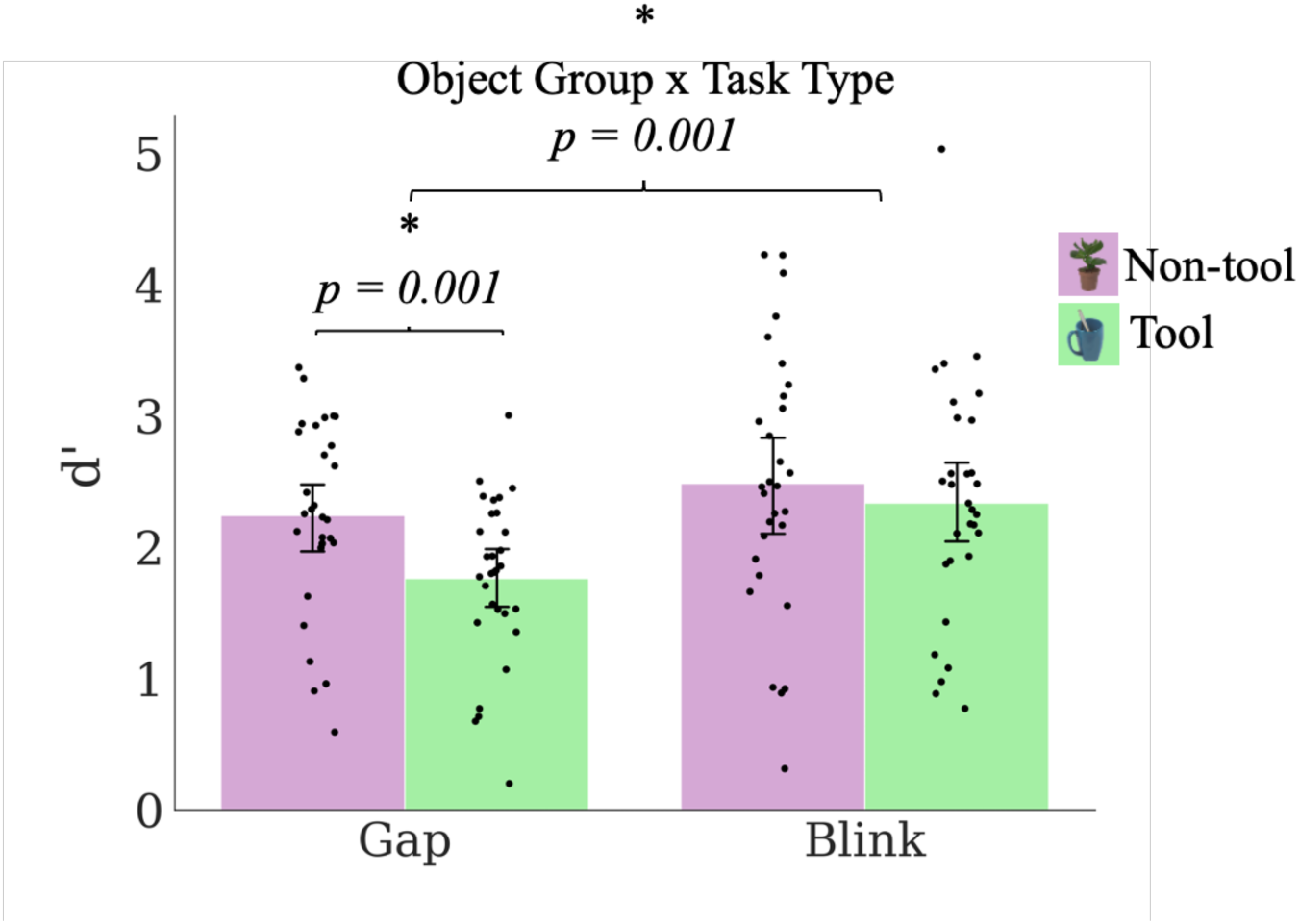
Experiment 3 results: a control manipulation of Experiment 1 in which real object images were used instead of line drawings.

### Experiment 4

While the first three experiments yielded strong supporting evidence that gap detection is better on non-manipulable objects than manipulable objects, perhaps resulting from bias toward the ventral pathway, manipulable objects failed to elicit an advantage in temporal resolution, failing to provide evidence of a dorsal pathway bias for manipulable objects. One explanation for the null effect is that the temporal gap detection task could also be construed as an abrupt onset detection task due to the object suddenly reappearing after the temporal blink. Abrupt onsets are known to be highly salient and easily detectable (Yantis and Jonides, 1984) and evoke strong activity in the lateral intraparietal area, a known attentional area with strong connections to the superior colliculus (Kusunoki et al., 2000). The strong connection between the LIP and the superior colliculus provides a mechanism by which abrupt onsets may bypass the parvo- and magno-cellular two visual pathways framework which we aimed to test, thereby collapsing any differences in the d’ observed for manipulable and non-manipulable objects. Another possible explanation for the null effect could be that the temporal gap detection as was too easy (ceiling effect), evidenced by high d’ values in the temporal task. In order to have a task that is more difficult and is more specifically targeted to the high temporal resolution of the magnocellular channel, the blink paradigm used in Experiments 1 through 3 was re-designed to a discrimination task^1^. Objects were presented for 80 ms, removed from the screen for either a 16 ms or 48 ms blink, and redisplayed for 48 ms (Fig. 4). Participants’ task was to indicate whether the blink duration was short or long, with d’ being calculated with short blinks being considered hits or misses and long blinks considered as correct rejections or false alarms. The prediction was that manipulable objects should have a higher d’ than non-manipulable objects due to the increased magnocellular input, and therefore temporal resolution, of the dorsal pathway. A t-test was used to analyze the difference between manipulable and non-manipulable objects in d’ (M_non-manipulable_= 1.562, 95% CI [1.354, 1.770]; M_manipulable_ = 1.620, 95% CI [1.385, 1.855]), with no significant effect being found (t(48) = -0.362, p = 0.719).

**Figure. 4.**
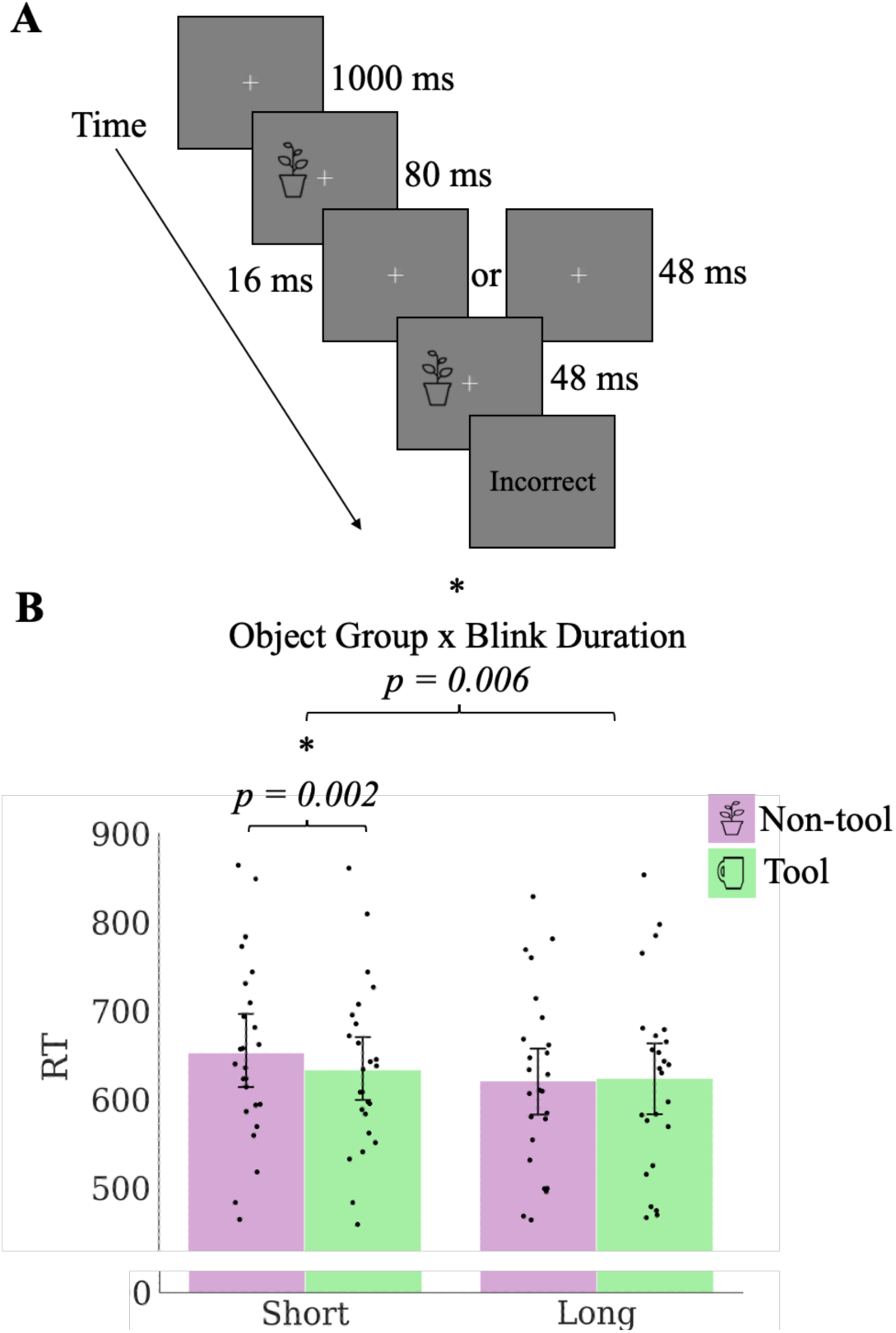
The paradigm and results for Experiment 4. A) Participants maintained fixation on the center cross. An object would then appear to the left or right of fixation. The objects would flicker by being removed from the screen for 16 ms or 48 ms. Participants were to indicate by keypress if the object flicker duration was short or long. B) The results from Experiment 4.

Due to the temporal nature of the task, RT was also analyzed, reasoning that the any benefit in temporal resolution for manipulable objects may manifest as a speeded reaction time rather than a sensitivity benefit. A two-way repeated-measures ANOVA was conducted on RT with object group (manipulable, non-manipulable) and blink duration (short, long) as within-subject variables. Only the RTs for correct responses were analyzed. The ANOVA revealed no significant main effect of object group (*F*(1, 24) = 4.080, *p* = 0.055, *η*_*p*_^*2*^ = 0.145) (Fig. 4). There was a significant main effect of blink duration (*F*(1, 24) = 4.850 *p* = 0.038 *η*_*p*_^*2*^ = 0.168) such that the short duration elicited higher RTs than did the long duration (M_short_ = 642.680, 95% CI [604.400, 680.960]; M_long_ = 621.997, 95% CI [581.794, 662.200]). The two-way interaction between object group and blink duration was significant (*F*(1, 24) = 9.011, *p* = 0.006, *η*_*p*_^*2*^ = 0.273) driven by a simple main effect in the short blink condition (*F*(1, 24) = 11.710, *p* = 0.002, *η*_*p*_^*2*^ = 0.328) such that manipulable objects had a lower RT for short blinks than did the non-manipulable objects (M_non-manipulable_ = 652.210, 95% CI [612.360, 692.060]; M_manipulable_ = 633.150, 95% CI [596.441, 669.859]). In accuracy, no effect was observed for object group (*F*(1, 24) = 3.666, *p* = 0.068, *η*_*p*_^*2*^ = 0.133), blink duration (*F*(1, 24) = 1.509, *p* = 0.231, *η*_*p*_^*2*^ = 0.059), nor the interaction (*F*(1, 24) = 2.000, *p* = 0.170, *η*_*p*_^*2*^ = 0.077), suggesting that all significant effects were absorbed by the response time measure and no speed-accuracy tradeoffs were observed, indicating consistency with response time results.

These results, specifically the simple main effect in the short gap condition with manipulable objects having a lower RT than non-manipulable objects, lend some support to our hypothesis that manipulable objects would elicit higher temporal resolution than non-manipulable object processing. While this result is not as strong as the observed benefit for non-manipulable objects in spatial resolution, nor is it manifested in sensitivity, but it does suggest that an advantage in temporal resolution is indeed present for manipulable objects.

Understanding the exact nature of this temporal resolution advantage may be a fruitful direction for future studies to take.

### Experiment 5

The last test of our hypotheses is derived from the neurophysiological differences between the dorsal and ventral pathways. It was reasoned that if the differences in spatial gap sensitivity for non-manipulable objects are mechanistically derived from the magnocellular and parvocellular dichotomy of the two visual pathways, then the effect should be modulated through manipulation of the processing within the pathways. Ambient red light has been shown to suppress activity in the magnocellular channel because of the large number of visually responsive cells with red inhibitory surrounds in their receptive fields (Wiesel & Hubel, 1966) and has been used to demonstrate the contribution of magnocellular processing in other behavioral effects such as fear processing and near hand effects (West et al., 2010; Abrams and Weidler, 2014). To test our hypothesis that the spatial resolution difference between manipulable and non-manipulable objects that we observed in Experiments 1 and 3 is due to the differential input of the magnocellular channel to the two visual streams, the spatial resolution paradigm from Experiment 1 was used with the manipulation of background color varying between green or red (Fig. 5A). Due to the suppression of the magnocellular channel by red light, it was predicted that the color of the background should modulate the perceptual difference in spatial gap detected observed in Experiment 1. A two-way repeated-measures ANOVA was conducted on d’ with object group (manipulable, non-manipulable) and background color (green, red) as within-subject variables. The ANOVA revealed a significant main effect of object group (F(1, 25) = 17.171, p < 0.001, *η*_*p*_^*2*^ = 0.407) with non-manipulable objects having a higher average d’ than manipulable objects (M_non-manipulable_ = 2.250, 95% CI [1.941, 2.559]; M_manipulable_ = 2.022, 95% CI [1.757, 2.287]) (Fig. 5B). Note that this main effect per se is a second replication of the spatial effect seen in Experiments 1 and 3. There was no significant main effect of background color (F(1, 25) = 2.138, p = 0.156, *η*_*p*_^*2*^ = 0.079) but, as predicted, there was a significant two-way interaction between object group and background color (F(1, 25) = 4.444, p = 0.045, *η*_*p*_^*2*^ = 0.151) such that the effect was increased with the red background. It was hypothesized that if the perceptual differences between manipulable and non-manipulable objects are mechanistically derived from the magno- and parvo-cellular processing in the two visual pathways, then suppression of the magnocellular processing with red light should modulate the effect. The results of Experiment 5 support our hypothesis by demonstrating an increase of the perceptual difference between manipulable and non-manipulable objects with red light.

**Figure 5.**
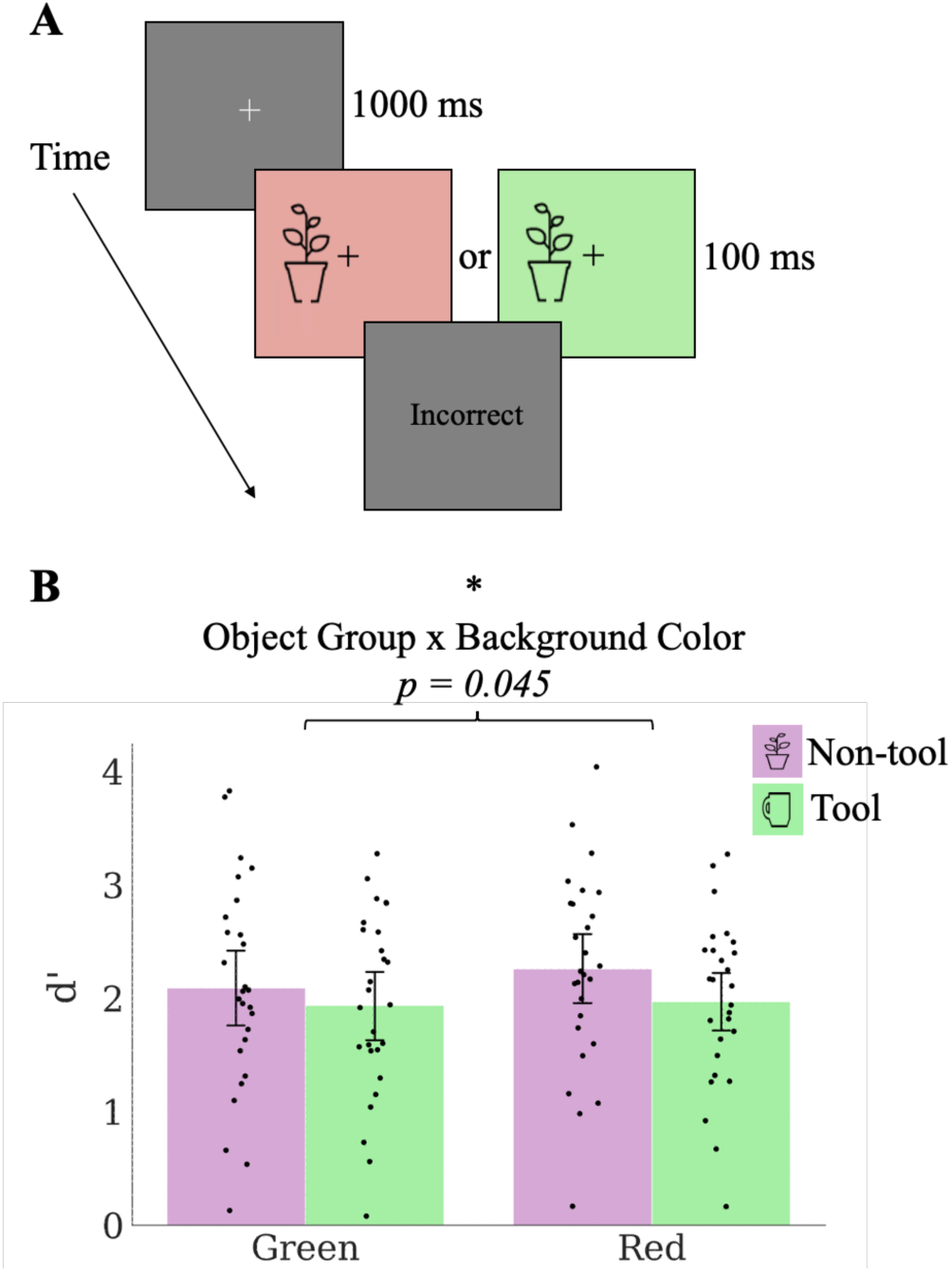
The paradigm and results of Experiment 5. (A) Participants maintained fixation on the center cross. An object then appeared to the left or right of fixation. As in Experiment 1, participants were asked to detect the presence of small gap in the bottom outline of the object. The background color (red or green) was manipulated. (B) The results from Experiment 5.

## Discussion

We hypothesized that manipulable and non-manipulable objects, due to differential recruitment of the visual pathways, would elicit perceptual differences stemming from the characteristic spatial and temporal resolutions associated with the magno- and parvo-cellular inputs. In five experiments we provide strong evidence in support of behavioral consequences driven by physiological and anatomical differences of the two visual pathways, and argue that semantic knowledge of object manipulability guides processing along a particular pathway. If an object has a strong action association, processing is largely determined by activity in the dorsal pathway which is not found with objects that lack strong action association (Chao & Martin, 2000; Noppeney et al., 2006; Mahon et al., 2007). The increased level of activity in the parietal regions endows the perception of action-associated objects with greater access to the magnocellular channel which preferentially courses through the dorsal pathway. Without the enhanced dorsal pathway activity evoked by action associations, object processing is more dependent on the ventral pathway, which has a higher ratio of parvocellular channel input than does the dorsal pathway (Maunsell et al., 1990; Baizer et al., 1991; Ferrera et al., 1992). Based on differential magno- and parvo-cellular input to the two pathways, we predicted that the perception of objects with strong action associations results in an increased temporal resolution and the perception of objects without such associations results in an increased spatial resolution. In the experiments reported here, we offer strong evidence that objects are perceived with different spatial resolutions, and some evidence that objects are perceived with different temporal resolutions, depending on object semantic knowledge of manipulability. In two follow-up control experiments, further evidence was provided in support of the hypothesis that the difference between manipulable and non-manipulable objects in the spatial task was driven by the semantic knowledge of the objects rather than possible low-level visual features. Namely, it was observed that the effect is curtailed by impeding the access of object semantic knowledge through inversion and that the perceptual differences between manipulable and non-manipulable objects replicates with a separate set of more realistic object images (with different low-level properties). Lastly, to test if the differing proportions of magno- and parvo-cellular input are responsible for the perceptual differences that were observed in our first four experiments, ambient red light was used to suppress activity of the magnocellular channel. It was observed that this increased the perceptual difference in the spatial task between manipulable and non-manipulable objects.

Based on the evidence provided, we argue that semantic knowledge of object manipulability, as defined by strong associations with an action appropriate for the item, generates the perceptual differences between manipulable and non-manipulable objects.

Previous studies have shown that manipulable objects evoke a larger degree of dorsal pathway activity than do non-manipulable objects (Chao & Martin, 2000; Almeida et al., 2013). However, the origin of this difference and consequent perceptual differences between manipulable and non-manipulable objects is poorly understood.

We propose that the differences we observe between manipulable and non-manipulable objects derive directly from the near-hand effect reported by Gozli et al. (2012) in which stimuli presented proximally to the participants’ hands evoked a benefit in spatial resolution, similar to the non-manipulable objects used in the experiments presented here, and stimuli presented distally from the participants’ hands evoked a benefit in temporal resolution. A possible mechanism supporting a connection between the near-hand effect and the manipulable vs. non-manipulable effect presented here derives from the bimodal cells in the anterior parietal cortex (Graziano & Gross, 1993). The bimodal cells have a somatosensory receptive field covering a part of the hand (a property possibly underlying the near-hand effect) and a visual receptive field corresponding to the visual field near the associated hand area. When an object is shown, if that object does not have a strong action association then the activity of the bimodal cells is unaffected. After the organism has learned to manually manipulate the object, the bimodal cells respond to visual presentation of the object even without somatosensory input (Zhou & Fuster, 2000). Thus, the bimodal cells, with their responsiveness to hand location and their ability to become visually activated, may be a driver of the dorsal pathway’s object selectivity (Vaziri-Pashkam & Xu, 2017; Kastner et al., 2017; Freud et al., 2016), leading to the dorsal pathway bias evoked by manipulable objects (Chao & Martin, 2000). The object-specific neural activity in the dorsal pathway, divorced from its need for somatosensory input, and largely derived from magnocellular input, could then be read out during an identification task, leading to heightened temporal precision for manipulable objects. Taken together, our results demonstrate that object semantic knowledge determines the processing bias of the object and evokes subsequent behavioral repercussions for perception and for action. This finding, in conjunction with the finding that manipulability has an influence on how attention interacts with object perception (Gomez et al., 2018), may point to manipulability, supported by a detailed neural mechanism, as an exemplar of cognitive penetrability. Additionally, our work underscores the need for careful consideration of object semantic knowledge, and its subsequent possible bias to either dorsal or ventral pathway, when object images are used not only in psychological research, but in applied settings such as display and product designs, environmental design, and in the design of various cognitive assistants.

## Author Contributions

D. Dubbelde and S. Shomstein designed the study. Testing, data collection, and data analysis were performed by D. Dubbelde under the supervision of S. Shomstein. D. Dubbelde and S. Shomstein wrote the manuscript. Both authors approved the final version of the manuscript for submission.

## Footnotes

1 We thank Ed Awh, from the University of Chicago, for the helpful suggestion.

